# SamQL: A Structured Query Language and filtering tool for the SAM/BAM file format

**DOI:** 10.1101/2021.02.03.429524

**Authors:** Christopher T Lee, Manolis Maragkakis

## Abstract

The Sequence Alignment/Map Format Specification (SAM) is one of the most widely used file formats in computational biology today and several tools have been developed to process it and use it. It is a flexible file format, used by many bioinformaticians on a daily basis. Despite its flexibility, SAM encoded files can often be difficult to query and understand. As genomic data are rapidly growing, structured and efficient queries on data encoded in SAM/BAM files is becoming critical. Importantly, any new tools should be able to support existing large datasets without requiring any data transformations. Here we introduce SamQL, an SQL-like query language for the SAM format with intuitive syntax that supports complex and efficient queries on top of SAM/BAM files and that can replace commonly used Bash one-liners employed by many bioinformaticians. SamQL is a complete query language that we envision as a step to a structured database engine for genomics. SamQL is written in Go, taking advantage of modern multicore compute systems and is freely available as standalone program and as an open-source library released under an MIT license, https://github.com/maragkakislab/samql/.

## Introduction

The advent of high-throughput sequencing has created an unprecedented availability of genomic datasets, highlighting the need for efficient storage and processing of large data. To address these requirements the Sequence Alignment Map (SAM) and its binary equivalent (BAM) file formats were developed (1). These file formats together with the, later developed, CRAM (2) format have been adopted by many bioinformatics software, including almost all alignment programs (3–6). Each record in the SAM format has several descriptive fields including alignment coordinates, sequence information, sequence and mapping quality and others.

Along with these file formats, SAMtools and other specialized programs (7–9) were developed to enable access to and processing of the encoded data. Nevertheless, despite these developments, advanced complex queries remain substantially difficult to perform and the web is flooded with related questions from investigators that usually end up using simple or not so simple Bash commands. Additionally, while SAMtools and other programs have a wide variety of functions and options, some of them can be unintuitive and require the user to often refer to the documentation leading to coding time increase and possibly inefficient or error-prone software.

These limitations are now becoming even more apparent due to the rapid increase in the size of genomic data (10) that is introducing new major challenges regarding access, security and data management (11). To address these limitations a new structured database engine and a supporting flexible query language that can allow organized data access is needed.

Here we introduce SamQL, a command line tool and library, that allows for SQL-like queries on top of the SAM/BAM format. SamQL has intuitive and flexible syntax and is parallelizable by taking advantage of modern multicore hardware architectures. SamQL can eventually enable the development of a large genomics database on top of SAM/BAM files, enabling complex queries for genomic datasets.

## Results

### Syntax and structure

SamQL aims to provide a user-friendly, flexible syntax and to support a parallelizable, database for genomics. SamQL was developed in the Go programming language that has been designed for multicore and large-scale network servers and big distributed systems. Internally, SamQL uses two robust and flexible bioinformatics libraries biogo (12) and hts (13) that provide a clean interface to common bioinformatics file formats. SamQL consists of a complete lexer that performs lexical analysis, and a parser, that together analyze the syntax of the provided query.

We designed SamQL queries to look very similar to SQL queries to make the system intuitive to use and to substantially reduce the learning curve. To support SAM-specific data extraction, SamQL recognizes SAM fields by their corresponding names defined in the specification (i.e. QNAME, FLAG, RNAME, POS, MAPQ, CIGAR, RNEXT, PNEXT, TLEN, SEQ, QUAL) and assigns them as language keywords. These keywords are dynamically replaced by the actual concrete values upon code execution. The SamQL model is very flexible and additional keywords can be added to support future requirements. For example, the LENGTH keyword has been added to correspond to the alignment length and is automatically evaluated for the user. Fig. 1A shows an example of the SamQL syntax and highlights the flexibility of the query system. Importantly, the query is evaluated once at the beginning of the search making the model lightweight and reducing computational time. To support this, SamQL builds an abstract syntax tree (AST) corresponding to the query. This is then parsed, depth-first, to progressively build a function closure that encapsulates the whole tree (Fig. 1B). The closure contains the entire filtering criteria, can accept a SAM record for filtering, and returns a boolean value indicating whether the record passes all criteria.

**Fig 1.**
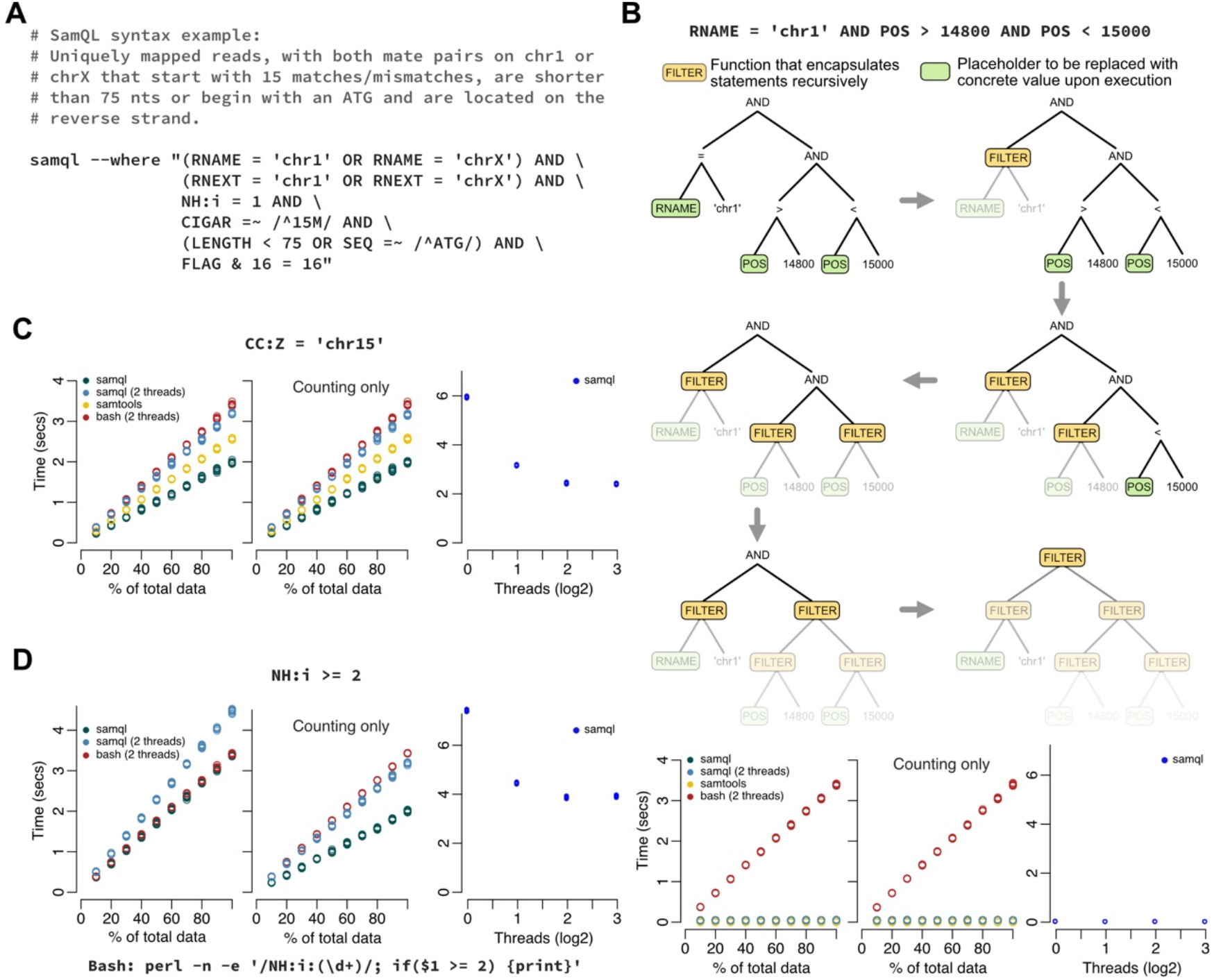
SamQL offers an intuitive and structured query language for SAM/BAM files. (**A**) Example of a complex SamQL query. SamQL is extremely flexible and can support searching on any fields of interest, including optional fields, with intuitive syntax similar to standard SQL. (**B**) Execution algorithm and performance of a query in SamQL. A range query is used as an example. Top: The abstract syntax tree constructed from the query. A FILTER function returns a single boolean value that encapsulates all filters below. The whole tree is progressively replaced by a single FILTER function that is used for record filtering. Bottom: From left to right, the plots correspond to runtime measurements for printing, counting, and multi-thread execution respectively. Colors correspond to SamQL: 1 thread (dark blue), 2 threads (blue), SAMtools (yellow), and naive Bash (red). (**C**) Same as B (bottom) for string query on tag “CC:Z” to be equal to “chr15”. (**D**) Same as B (bottom) for integer query on tag NH:i to be greater than or equal to 2. SAMtools does not support this query. Perl is used instead of Awk and the command is shown below.

### Performance

While our primary focus in SamQL was syntax flexibility, we wished to also test the computational performance of SamQL against other widely used tools. Therefore, we compared it against SAMtools and a naive Bash implementation. We tested on three different queries, each one with different levels of syntax complexity and computational requirements. We repeated each query 10 times on varying input sizes to validate the accuracy of our measurements. All comparisons were run on an Intel Xeon, 48 cores, 384Gb memory compute server. We used a publicly available BAM file with 3,363,576 records. To evaluate scalability, we randomly sampled the input BAM file into sizes of 10% increments and performed measurements on all subsets. To evaluate the query performance fairly and decouple it from IO we measured the execution time both when printing to an output file but also just counting the filtered reads (without outputting). Finally, we also measured the capabilities of SamQL to parallelize execution and use multiple cores.

As a first test, we decided to compare performance on filtering against the CC:Z tag that uses string matching and is supported by all methods. Filtering on such optional tags is a great test because it forces all methods to read the entire SAM record thus decoupling optimizations that depend on skipping optional SAM fields. We find that SamQL, while being infinitely more flexible in its syntax, performs on par with the other methods even when bound to only use two threads, an input/output (IO) and a compute (Fig. 1C). When no thread bound is enforced, SamQL performs better than both naive Bash and SAMtools. Importantly and as expected by design, execution time increases linearly with the input size, that is a feature that allows SamQL to be used for very large datasets.

To highlight the power of the syntax flexibility, we then wished to filter on the NH:i tag that involves numerical comparisons. While this is an intuitive and straightforward query change in SamQL (Fig. 1D, top), it raises significant issues for the other methods. First, SAMtools does not even support numerical comparison in the query, while the naive Bash implementation becomes substantially more complex (Fig. 1D, bottom). Again, we find that SamQL execution time is on par or faster than Bash and that execution time again increases linearly with time (Fig. 1D). Interestingly, we find that when many records are involved, as in this case, the improved SamQL performance compared to Bash, is even higher when counting records (Fig 1D, middle panel) compared to outputting (Fig 1D, left panel) indicating a very performant SamQL query execution that is possibly bound by the external, underlying Go libraries and that would benefit by further improvement in those.

Finally, we evaluated the performance for a reasonably complex range query. Range queries on genomic or transcriptomic coordinates are very commonly included in bioinformatics analysis. Therefore, BAM files are usually indexed to achieve fast retrieval of alignments overlapping a specified region(14). SamQL, in contrast to SAMtools that requires indexing and runs only on BAM files, can execute range queries on indexed and not indexed BAM or SAM files, albeit much faster when indexing exists. Our data show that SamQL performs on par with SAMtools and orders of magnitude faster than a naive approach, as expected (Fig 1B). Overall, all out tests indicate that SamQL offers infinite syntax flexibility while maintaining performance at a level comparable or faster than other tools and is able to take advantage of parallel computing.

## Discussion

SamQL enables complex queries in a straightforward and intuitive way. It is simple to use and can replace the majority of one-liners used by bioinformaticians, thus helping to reduce errors. While SamQL is able to utilize and benefit from multicore systems we find that performance improvement plateaus at approximately 4 threads indicating that there is room for optimization improvements in the underlying libraries. Importantly, SamQL is a lightweight library that can act as a query language for future development of a complete, large genomic database. We envision a database running on top of existing SAM/BAM files where investigators would be able to easily search through every file source in the database for reads of interest.

## Funding

This research was supported by the Intramural Research Program of the National Institute on Aging, National Institutes of Health.

